# SpatialDiffusion: Predicting Spatial Transcriptomics with Denoising Diffusion Probabilistic Models

**DOI:** 10.1101/2024.05.21.595094

**Authors:** Sumeer Ahmad Khan, Vincenzo Lagani, Robert Lehmann, Narsis A. Kiani, David Gomez-Cabrero, Jesper Tegner

**Affiliations:** Biological and Environmental Science and Engineering Division, King Abdullah University of Science and Technology (KAUST), Thuwal 23955-6900, Saudi Arabia; SDAIA-KAUST Center of Excellence in Data Science and Artificial Intelligence, Thuwal 23952, Saudi Arabia; Institute of Chemical Biology, Ilia State University, 0162 Tbilisi, Georgia; Algorithmic Dynamic Lab, Department of Oncology and Pathology, Karolinska Institute, Stockholm, Sweden; Unit of Computational Medicine, Department of Medicine, Center for Molecular Medicine, Karolinska Institutet, Karolinska University Hospital, L8:05, SE-171 76, Stockholm, Sweden; Translational Bioinformatics Unit, Navarrabiomed, Universidad Pública de Navarra (UPNA), IdiSNA, Pamplona, Spain; Computer, Electrical and Mathematical Sciences and Engineering Division, King Abdullah University of Science and Technology (KAUST), Thuwal 23955-6900, Saudi Arabia; Science for Life Laboratory, Tomtebodavagen 23A, SE-17165, Solna, Sweden

## Abstract

Spatial Transcriptomics (ST) allows deep characterization of the 2D organization of expression data within tissue slices. The ST technology provides a tissue contextualization of deep single-cell profiles. Recently, numerous computational and machine learning methods have addressed challenges such as data quality, augmentation, annotation, and the development of integrative platforms for data analysis. In contrast, here we ask whether <u>unseen spatial transcriptomics data can be predicted and if we can interpolate novel transcriptomic slices. To this end</u>, we adopt a denoising diffusion probabilistic-based model (DDPM) to <u>demonstrate the learning of</u> generative ST models for several tissues. Furthermore, our generative diffusion model interpolates (predicts) unseen slices located “between” the collected finite number of ST slices. This methodology set the stage for learning predictive deep 3D models of tissues from a finite number of spatial transcriptomics slices, thus harboring the advent of AI-augmented spatial transcriptomics.

---

The twin rise of single-cell genomics, producing high-resolution atlases of different cells, and spatial transcriptomics (ST), uncovering the cellular transcriptional organization within tissues, impacts fundamental cell biology and translational research^1^. Recently, a wide range of computational and machine learning methods target challenges such as data quality and augmentation (imputation, normalization), annotation (deconvolution of cell types, clustering, resolution), and tissue analysis (spatial gradients, extracting hierarchical organization), and development of integrative platforms (software, databases, data management)^2–4^.

While the bulk of the work has been 2D Spatial Transcriptomics at different length scales^4^, increasing efforts are homing to the potential of 3D Spatial Transcriptomics. Recently, progress has been achieved in aligning ST slices using probabilistic models^5^ or aligning slices using a 3D neighborhood graph^6^ into a local coordinate system^7,3^.

Here, we take the next logical step, learning a generative ST model for any tissue. Specifically, for a given finite set of ST slices, we ask if we can predict or interpolate unseen ST slices beyond a finite set of ST slices (Figure 1a). To this end, we first assessed whether a denoising diffusion probabilistic model^8^ (DDPM) could serve as a generative model for ST. The trilemma – the balancing act between quality, diversity, and speed - of generative models^9^ means that both variational encoders and normalizing flows suffer from lower quality. In contrast, Generative Adversarial Networks are prone to mode collapse, thus rendering them less suitable for producing diversity. Recently, the DDPM model^8^ has achieved a faster subsampling, becoming a practical model for learning via a forward noise process and neural network learning the reverse diffusion process (Figure 1b). We hypothesized that DDPM models are a suitable candidate for the reconstruction of ST slices as DDPM has recently achieved state-of-the-art in image generation (DALLE 2), super-resolution, image denoising, and inpainting^10^.

**Figure 1:**
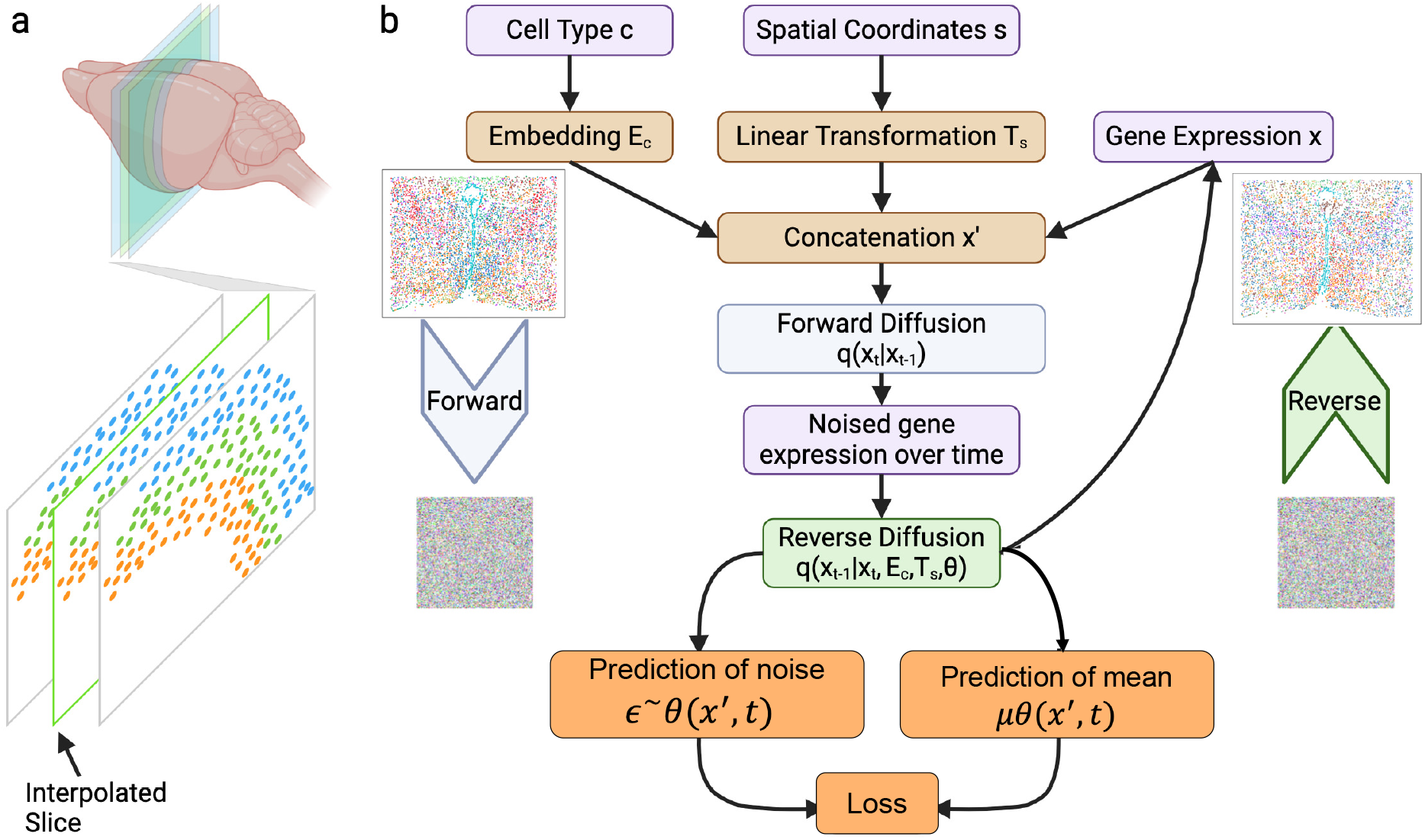
Schematic of stDiffusion: **a)** depiction of interpolation across slices of brain **b)** stDiffusion for in silico generation and interpolation of spatial transcriptomics data. Details in methods section.

To evaluate whether DDPM models are suitable for the reconstruction of ST slices, we trained a DDPM model (stDiffusion) (Figure 1b, Methods) on mouse MERFISH^11^ slice data. For a given slice, the input to the model is gene expression for each spot, along with the cell type and the spatial coordinates, encoded via the embedding and linear layers and activation functions (Methods). The DDPM model uses concatenated features to predict the original data from its noised version. We use spot-wise clustering to compare the structure of ground truth and generate data using the neighborhood enrichment test^12^ to identify spatially enriched clusters. Visualizing the clustering (Supp Fig S1a,b) demonstrates the model’s ability to closely replicate the structure and relationships found in the reference data (Figure 1b). Notably, embedding contextual features such as cell type and spatial coordinates and concatenating them with the gene expression data enhances the model*’*s predictive accuracy during reverse diffusion. Thus, as a proof-of-principle, we provide the first demonstration that a DDPM model can generate and reconstruct spatial transcriptomics data.

Next, we asked whether stDiffusion could generate unseen ST slices by interpolating within a given slice and between two adjacent slices (Figure 1a). The idea is to harness the DDPM’s ability to learn the underlying data distribution through gradual transformations from noise to data, thus effectively navigating a learned latent space. This task corresponds to smooth interpolations between observed data points. To this end, we trained stDiffusion on a subset of ST slices and then tested if non-border held-out slices could be predicted. We used three different datasets originating from various tissues and technologies.

In brief, we extract the gene expression data and spatial coordinates for a given layer and Bregma values for each slice, which serve as reference points. These coordinates are normalized to ensure that distance calculations between points are consistent and meaningful across the dataset’s varying scales of spatial dimensions. Stochastic gene expression encoding is essential for preparing the data for interpolation in the model’s latent space, allowing us to simulate the variability inherent in biological data. For each spot in a given target slice, we use Euclidean distance to identify the closest points in the adjacent reference slices, providing a basis for choosing the most relevant features for interpolation (Methods). The interpolation process leverages a predetermined factor (λ) to blend the noised feature representations of the closest points from the reference slices. This factor can be adjusted to explore different blending ratios, allowing for a controlled simulation and generation of a continuous transition between slices (Supp Fig 3, Fig 4, and Fig 5). The proximity-based selection of features, informed by the calculated distances, ensures that the interpolated data respects the spatial continuity and variation observed in the original slices. Following the interpolation in latent space, we employed the reverse diffusion process to reconstruct the interpolated gene expression profiles from their distorted versions. As a baseline for comparing the performance of the DDPM model, we average the expression profiles of two reference slices, which correspond to a linear interpolation of the two nearby slices (Methods).

Using this workflow, we first challenged stDiffusion to interpolate within a slice of the human dorsolateral prefrontal cortex (DLPFC^13^, Methods) between layers. Using sample profiling layers Layer_1 through Layer_6 and white matter (WM), we aimed to predict the Layer_4 ST between Layer_3 and Layer_5 (Figure 2 a-c). stDiffusion performs remarkably well (Figure 2b,c) in interpolating layers in a slice that were not included during the initial and validation phases while preserving the spatial neighborhoods within these layers compared to linear interpolation (Figure 2b,c). Large neighborhood enrichment values along the diagonal (Figure 2b) indicate a robust within-cluster relationship, demonstrating cells of the same type are located near each other.

**Figure 2:**
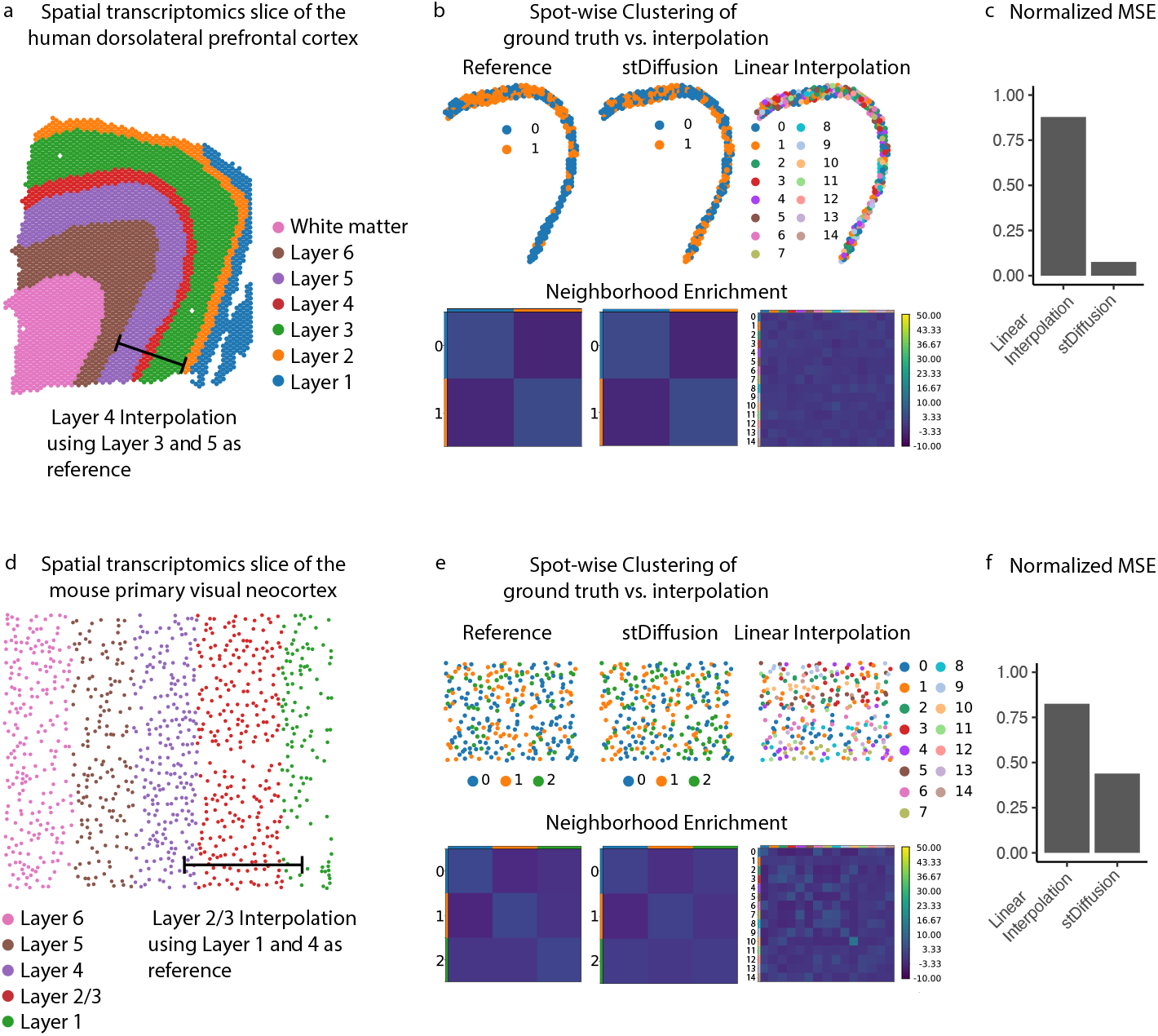
Interpolation across human dorsolateral prefrontal cortex (DLPFC) with stDiffusion: **a)** visualization of DLPFC slice layers data in spatial coordinates, **b)** shows Leiden clustering in spatial coordinates and neighborhood enrichment heatmap for the reference DLPFC ground truth: Layer_4), this layer exhibits two distinct clusters. This clustering pattern is effectively mirrored by stDiffusion, which also identifies two clusters aligning closely with the spatial organization observed in the reference data compared to linear interpolation. The intense colors in the heatmaps along the diagonal indicate a strong within-cluster relationship, demonstrating cells of the same type are located near each other. The off-diagonal element shows the enrichment score between the two clusters, indicating their proximity relationship. The consistency in the intensity both along the diagonal and off-diagonal elements suggests the stDiffusion model effectively captures the spatial interactions of the DLPFC’s cellular composition as observed in the actual data, **c)** normalized mean square errors between the ground truth and interpolated slice, **d)** visualization of Starmap data in spatial coordinates, **e)** shows Leiden clustering in spatial coordinates and neighborhood enrichment heatmap for the reference (ground truth: L2/3) layer. Interpolation results with stDiffusion show its efficacy in interpolating and capturing the spatial structure observed in the ground truth. In contrast, results of linear interpolation show its limitations in interpolating and capturing the spatial structure observed in the ground truth, **f**) normalized mean square errors between the ground truth Starmap and interpolated Starmap slice, depicting better performance of stDiffusion as compared to linear interpolation.

The pattern is very similar in the reference and the interpolated data, suggesting that the stDiffusion model has effectively recapitulated the actual spatial relationships in the real tissue data. stDiffusion produced a robust positive association between interpolated and actual expression values (Supp Fig 2a), indicating that the spatial arrangement of gene expressions in the interpolated layer closely resembles that in the ground truth layers.

Next, we asked whether stDiffusion could navigate across the layers within a single slice of the mouse visual cortex. Here we used the Starmap^14^ encompassing distinct cortical layers L1, L2/L3, L4, L5, and L6. The task is to interpolate the intermediate layer L2/L3 by interpolating between layers L1 and L4 (Figure 2 d-f), thus recapitulating the organization of clusters as observed in the real data. To this end, stDiffusion accurately reconstructs the omitted layer L2/L3 (Figure 2e). Notably, stDiffusion outperforms a linear interpolation method (Figure 2e,f), which falls short in capturing the spatial distribution and cluster neighborhood structure (Figure 2e, Supp Fig 2d). stDiffusion effectively (Supp Fig 2c) interpolates and generates gene expression profiles of the target layer.

Finally, we used the mouse MERFISH^11^ data of 12 consecutive slices of the hypothalamus preoptic region (Figure 3a-d). The results (Figure 3b,c,d) show that stDiffusion effectively interpolates between slices, successfully preserving the structure of cluster neighborhoods as seen in the original data. For instance, in the MERFISH data slice, the spatial structure of cluster neighborhoods, such as those of cluster 8 and cluster 11, is maintained in the interpolated slice, aligning with observations in the ground truth slice (Figure 3d). The superior performance of stDiffusion is in stark contrast with that of linear interpolation, which fails to preserve the cluster neighborhood structure (Figure 3d). Furthermore, we use Leiden clustering to identify spatially distributed clusters and compare the structure of ground truth and generated data. Visualizing clustering’s in spatial coordinates (Figure 3b) demonstrates the model’s ability to closely replicate the structure and relationships found in the reference data (Figure 3a). We also conducted a normalized Spearman correlation analysis between the expression levels of all genes in the ground truth slice and those in the interpolated slice for each spot. These correlation values were then plotted at their respective spatial coordinates. The findings (Supp Fig 2e) indicate that stDiffusion accurately maintains the spatial distribution of gene expression levels in the interpolated slice, consistent with the original data.

**Figure 3:**
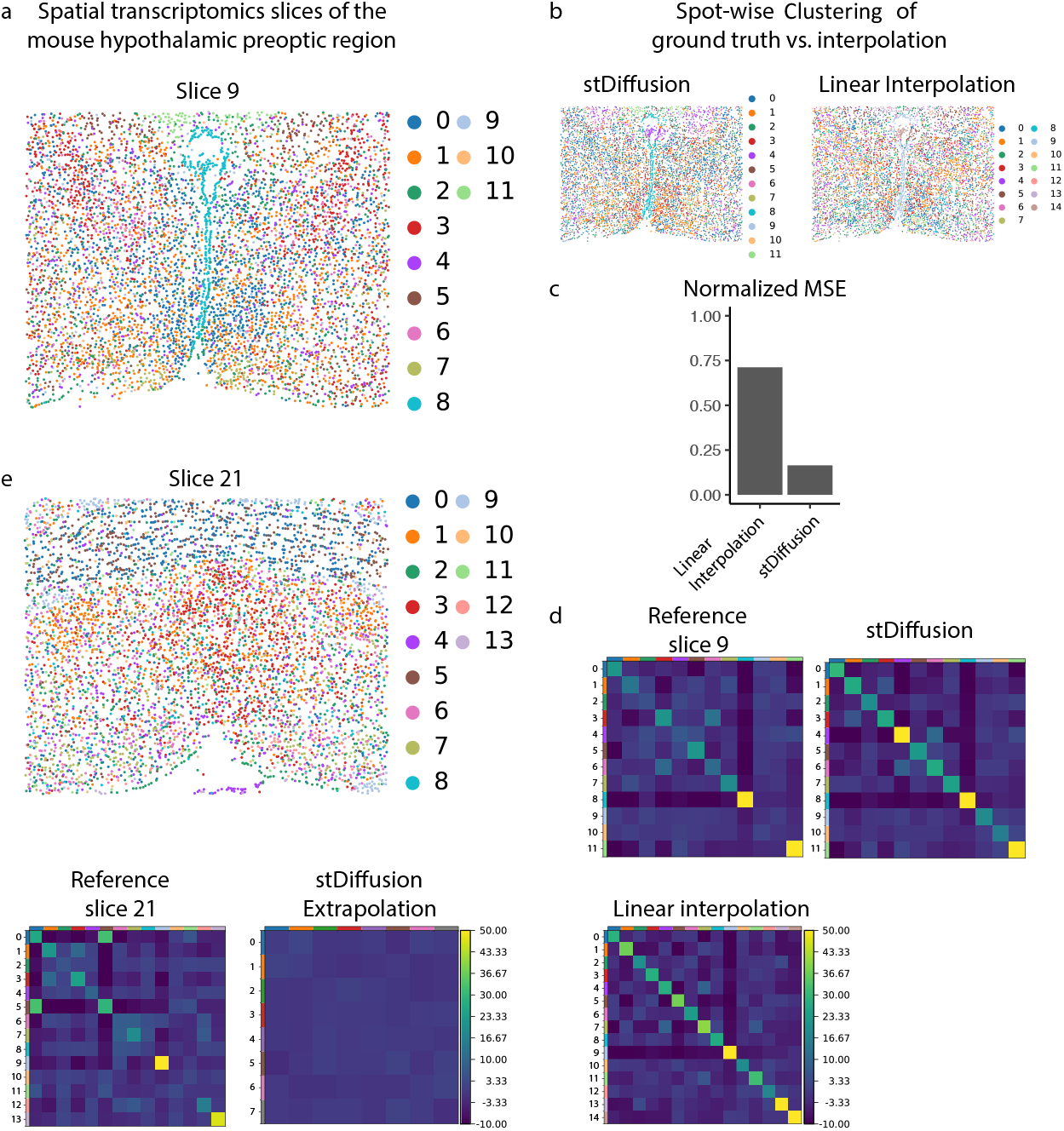
Interpolation and extrapolation - MERFISH slices with stDiffusion: **a)** visualization of ground truth slice (Bregma 9 and Bregma 21) data in spatial coordinates, **b)** Leiden clustering results of interpolated slice in spatial coordinates, **c)** normalized mean square errors between the ground truth MERFISH slice and interpolated slice, depicting better performance of stDiffusion as compared to linear interpolation, **d)** neighborhood enrichment for reference (ground truth: slice (Bregma 9)). Results of interpolation with SpatialDiffusion showing its efficacy in interpolating and capturing the spatial structure, e.g., cluster 8 cells observed in ground truth MERFISH data reconstructed well in the interpolated slice, whereas heatmap of neighborhood enrichment from linear interpolations showing its inefficiency in retaining the spatial structure, **e)** neighborhood enrichment heatmap of the ground truth slice for extrapolation. The neighborhood enrichment heatmap shows the limits of stDiffusion when performing extrapolation (out-of-distribution prediction).

In summary, stDiffusion can learn ST data from a single slice and predict held-out slices, thus effectively interpolating between a finite set of ST slices. Given that the DDPM model can solve an in-distribution challenge, we next asked whether the stDiffusion model could extrapolate outside the collected spatial region. Since such an out-of-distribution will not necessarily have a similar statistical distribution as the collected data, we would expect, as a control, stDiffusion to have a lower performance. To this end, we trained the stDiffusion model using all available data slices from the posterior region of the mouse brain while excluding slice 21 (Figure 3e upper) from the anterior region during training and validation. Our findings validate that stDiffusion’s performance in extrapolation scenarios, as depicted in (Figure 3e lower) is not as robust as in interpolation. This finding serves as a controlled experiment delineating the predictive capabilities of a DDPM model. The extent to which extrapolation is possible will naturally depend on the distributional similarity between the training and testing datasets. Here, there is no reason to expect the posterior region of the mouse brain to incorporate sufficient information to predict the spatial transcriptomics structure in the anterior region. This suggests an experimental sampling strategy such that different areas are sampled since our results demonstrated that the stDiffusion model could interpolate and predict data within regions where samples have been collected.

In our study, we utilized the novel stDiffusion model based on DDPM^15^ to address two pivotal challenges in spatial transcriptomics: the generation of in silico data and the interpolation of gene expression across tissue slices and layers.

Extensive work has been carried out on what could be referred to as “local” challenges of 2D transcriptomics. This includes improving data quality, normalization, deconvolution of cell types, clustering, spatial gradients, and efforts to extract a hierarchical organization from 2D spatial transcriptomics data^2–4^. While some recent studies^5^,^6^,^7^ have begun to address how to align 2D slices in a 3D coordinate system, those studies still suffer from a finite number of samples, which by necessity means that the physical gaps between the slices remain. Here, we take the next logical step: to learn generative models from data and use them as predictive models for ST data. Our demonstration of the predictive power for interpolation sets the stage for building generative 3D models of tissues, thus transforming a set of discrete slices into a continuous generative 3D representation of tissues. The journey towards a generative 3D spatial transcriptomics model entails robust segmentation, a challenge that might be well suited for a DDPM type of model^10,16^. A generative 3D Spatial Transcriptomics AI program^16^ combining multimodal cellular analysis and tissue localization may impact fundamental biology, computational modeling of organs^17^, and precision medicine^18^.

## Methods

### Data Preprocessing

We developed and evaluated SpatialDiffusion (stDiffusion) using three different datasets. The MERFISH^11^ dataset from^12^ was used to assess the in-silico generation and interpolation performance of stDiffusion across distinct slices. MERFISH consists of 12 consecutive slices from the mouse hypothalamic preoptic region. The dataset is segmented based on the Bregma reference point, allowing for the exclusion or inclusion of specific slices to tailor our training, validation, and test sets for focused analyses. Each instance comprises gene expression values, classified cell types, and two-dimensional spatial coordinates, facilitating a multifaceted approach to understanding spatial gene expression patterns. The dataset has 73655 spots and 161 genes per spot. The second dataset we used for interpolation across layers is the mouse visual cortex Starmap^14^ data obtained from^19^ in the preprocessed format with 984 spots and 1020 genes per spot. This dataset has spots of gene expression profiles for L1, L2/L3, L4, L5, and L6 layers. The third dataset is the human dorsolateral prefrontal cortex from (DLPFC)^13,19^. Specifically, the DLPFC dataset includes 12 human DLPFC sections sampled from three individual experiments^13^. The number of spots ranges from 3498 to 4789 for each section, and the original authors have manually annotated the area of DLPFC layers and white matter (WM). We used one sample from this dataset, and we preprocessed with the SCANPY^20^ package, and used 3000 highly variable genes with 3431 spots as an input to our stDiffusion model. Additional information, such as spatial coordinates and layers, is included in the dataset. For DLPFC dataset, we applied our stDiffusion model to interpolate across layers, i.e., Layer_1, Layer_2, Layer_3, Layer_4, Layer_5, Layer_6, and white matter (WM).

### The SpatialDiffusion Model

In spatial transcriptomics, stDiffusion (Figure 1b) adapts Denoising Diffusion Probabilistic Models^8, 15^ (DDPM) principles to address the unique challenges posed by spatially resolved gene expression data. The model begins by learning the distribution of gene expression profiles across spatial coordinates, effectively capturing the spatial transcriptional landscape of the tissue. Through forward and reverse diffusion steps, stDiffusion interpolates missing gene expression data within this landscape, providing a continuous view of transcriptional activity across the tissue.

### In silico generation

The in-silico generation of spatial transcriptomics data aims at augmenting existing datasets with synthetic yet biologically plausible data points. This approach enhances the dataset’s density, providing a more continuous spatial representation of gene expression patterns. We trained the stDiffusion model on the preprocessed spatial transcriptomics data, allowing it to learn the underlying distribution of gene expression across spatial coordinates.

### Network Architecture

#### Diffusion Model

The core of our approach is a denoising diffusion probabilistic model (DDPM)^8^ specifically designed to handle the complexities of spatial transcriptomics data. This model incorporates an embedding layer for cell types and a linear transformation for spatial coordinates, ensuring both are integral to the learning process (Figure 1b). An embedding layer for cell type classification allows the model to interpret cell types as dense vectors of a specified dimension. A linear transformation is applied to spatial coordinates, mapping them to a similarly dimensioned space as the cell-type embeddings. A concatenation of gene expression data with the transformed cell type and spatial information is followed by a sequential network comprising linear layers and activation functions.

#### Enhancement of Model Inputs

Let *x* denote the gene expression data for a given sample, *c* denotes the cell type, and *s* represents the spatial coordinates. The cell type *c* is transformed through an embedding layer, and the spatial coordinates *s* are processed through a linear transformation to ensure they are in a compatible format for concatenation:

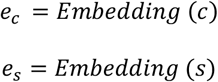

The enhanced input *x*^′^ to the model is then a concatenation of the original gene expression data with the transformed cell type and spatial information:

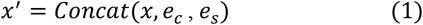

#### Forward Process (Diffusion)

The forward diffusion process gradually adds Gaussian noise to the data across several discrete time steps. The process at a specific time step t can be mathematically represented as follows:

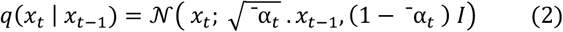

where:
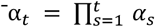 is the cumulative product of *α*_*t*_ = 1 − *β*_*t*_ with *β*_*t*_ being the variance of the noise added at step *t. I* is the identity matrix, ensuring the noise is independently added to each dimension.

#### Reverse Process (Diffusion)

During the reverse process, the enhanced input *x*^′^is used to predict the original data from its noised version. The model utilizes the concatenated features to predict the mean (*μ*) of the reverse process distribution:

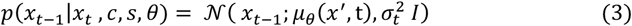

Where *μ*_*θ*_(*x*^′^, *t*) depends on the enhanced input *x*^′^, incorporating embeddings for cell type and spatial coordinates, along with the noised data *x*_*t*_.

Incorporating cell type and spatial coordinates into the stDiffusion allows the model to make more informed predictions during the reverse diffusion process. By embedding these contextual features and concatenating them with the gene expression data, the model can leverage the full spectrum of biological information available in the dataset. This procedure enhances the model’s predictive accuracy and the biological relevance of the reconstructed data, facilitating a deeper understanding of the spatial patterns of gene expression within tissue samples.

#### Loss function

The loss function aims to minimize the difference between the actual noise introduced during the forward diffusion process and the noise predicted by the model during reverse diffusion, incorporating cell type (c) and spatial information (s) as conditioning variables. To this end we use the Mean Squared Error (MSE) loss between the actual and predicted noise:

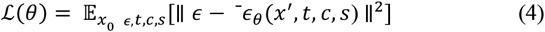

Where *ϵ* is the actual noise added to the original data *x*_0_ to obtain the noised version *x*_*t*_, ^-^*ϵ*_*θ*_(*x*^′^, *t, c, s*) is the noise predicted by the model and *x*_0_ is the original data, *t* is a randomly chosen diffusion time step, and *θ* denotes the model parameters. This objective encourages the model to accurately predict the noise at any given time step, thereby learning the reverse of the diffusion process to reconstruct the original data from noised observations.

### Interpolation with stDiffusion

#### Stochastic Encoding in the Forward Diffusion Process

The stochastic encoding process incrementally introduces Gaussian noise into the original gene expression data *x*_0_ across a series of discrete time steps. This stochastic encoding process transforms the data into a latent space where the original high-dimensional structure is preserved amidst the added noise. The process at a specific time step *t* can be mathematically represented as follows:

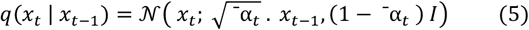

where:
^-^*α*_*t*_ is the cumulative product of 1 − *β*_*t*_ over time, indicating the proportion of the original signal preserved. *I* is the identity matrix, ensuring that noise is added isotropically.

#### Spatial Proximity Calculation

The interpolation between two slices involves calculating the spatial proximity to determine the blending factor *λ*. This is achieved by normalizing the coordinates and calculating distances to the target coordinates to ensure comparability:

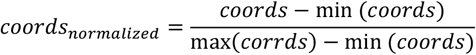

#### Distance Calculation

The Euclidean distance from each point in the slices to the target coordinates is calculated. The blending factor *λ* for each target point is then determined based on the proximity of points in *slice1* and *slice2* to the target coordinates. Our experimentation with *λ* values ranging from 0.1 to 0.9 has revealed its significant impact on interpolation quality, indicating that the optimized *λ* can vary across different datasets (Supp Fig3, Fig4, Fig5).

#### Latent Space Blending for Interpolation

Given noised data *x*_*t*,1_ from *slice1* and *x*_*t*,2_ from *slice2* at time *t*, the interpolation for a target spatial coordinate is computed by blending these representations based on spatial proximity. The interpolated data *x*_*t*,*interp*_ at time *t* is calculated as:

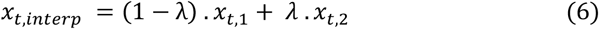

*λ* is adjusted based on the spatial proximity of the points in the slices to the target coordinate, influencing their contribution to the interpolated output.

#### Reverse Diffusion Process (Reconstruction)

The reverse diffusion process reconstructs the original data from the noised state to generate the interpolated gene expression data.

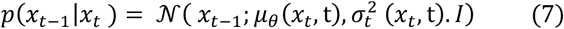

where:
*x*_*t* −1_ and *x*_*t*_ represents the gene expression data t two consecutive time steps, with *(t-1)* being closer to the original data.

*μ*_*θ*_(*x*_*t*_, *t*) and 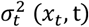 are the functions modeled by the neural network parameterized by *θ*. These functions predict the mean and variance of the Gaussian distribution from which *x* _*t* −1_ is sampled, given *x*_*t*_.

#### Loss function for interpolation

The loss function aims to minimize the difference between the actual noise added to the data during the forward process and the noise predicted by the model during the reverse process. For interpolation, we used mean squared error (MSE) between the actual noise *ϵ* and the estimated noise ^-^*ϵ*_*θ*_ given the noised data *x*_*t*_ and model parameters *θ*:

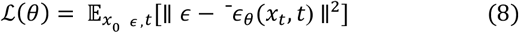

where:
*ϵ* is the actual noise vector sampled from a Gaussian distribution during the forward process, ^-^*ϵ*_*θ*_(*x*_*t*_, *t*)is the noise estimated by the model, given the noised data *x*_*t*_ at time *t* and 𝔼 denotes the expectation over the distribution of original data *x*_0_, noise *ϵ*, and time steps *t*.

For interpolation, the focus is on blending noised representations from different slices based on spatial proximity and then reconstructing the interpolated data through the reverse process. Although the loss function remains centered on noise prediction accuracy, the application in interpolation emphasizes the model’s capability to handle blended data from multiple sources and generate coherent interpolated outputs.

### Linear Interpolation

The linear interpolation method calculates the gene expression profile for a target slice by averaging the expression profiles of two reference slices. For each spatial coordinate in the target slice *S*_*2*_, the closest points in reference slices *S*_*1*_ and *S*_*3*_ are identified based on Euclidean distance. The gene expression profile for a point in target slice *S*_*2*_ is computed as the average of the gene expression profiles of the closest point in slices *S*_*1*_ and *S*_*3*_.

### Training Parameters

For tasks such as in silico generation and interpolation of spatial transcriptomics data, training parameters play an essential role in the performance of stDiffusion. The key parameters optimized for stDiffusion for the task of in silico generation and interpolation across slices and layers are noise schedule (*β*_*t*_) - linearly spaced from *1e-4 to 0*.*02*. The noise schedule directly impacts the model’s ability to learn the data distribution through diffusion, requiring tuning to match the complexity of spatial transcriptomics data. We used AdamW^21^ as an optimizer and learning rate (*lr = 1e-3)*. In addition to this stDiffusion is trained for 300 epochs with an early stopping criteria. To adapt the learning rate, using OneCycleLR promotes faster convergence and mitigates the risk of getting stuck in local minima. For interpolation tasks, most hyperparameters remain the same as those for in silico generation, reflecting the shared underlying model architecture and training strategy. However, specific considerations for interpolation include the interpolation Factor (λ), which ranges from 0.1 to 1.0 in increments of 0.1 in experiments. This factor controls the blend between the gene expression profiles of two slices.

### Evaluation Metrics

#### Neighborhood Enrichment Test

We used Squidpy’s ^12^ Neighborhood Enrichment Test. This test evaluates the spatial relationships between different clusters or types of cells within a tissue or dataset. It helps to understand how certain groups of cells are distributed to one another and whether there are significant patterns of co-localization or segregation. The test first identifies pairs of nodes (cells) belonging to specific classes or clusters (*i* and *j*). The next step is to count the sum of nodes that are proximal to each other, represented as *x*_*i*,*j*_. This count reflects the direct interactions or the degree of proximity between cells of two different clusters. To determine whether the observed proximity of cluster pairs is significant, the test compares the cluster distance against what would be expected by chance. The randomized background is computed by scrambling the cluster labels while keeping the spatial connectivity unchanged, effectively randomizing the distribution of clusters. This process is repeated multiple times (default:1,000 iterations) for a robust statistical comparison. From these iterations, the test calculates the expected means (*μ*_*i*,*j*_) and standard deviations (*σ*_*i*,*j*_) for the proximity counts of each cluster pair in the randomized configurations. A *z-score* is then computed for each cluster pair using the formula:

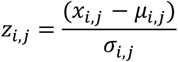

This *z-score* provides insights into the spatial organization of the tissue or dataset and helps to identify which cell types or clusters are more likely to be found near each other, potentially indicating functional relationships, shared microenvironments, or developmental pathways. To assess our model’s performance, we applied the Leiden clustering method to group spots/cells into clusters and used these clusters as the basis for our neighborhood enrichment test. This approach allowed us to directly compare the spatial relationships observed in both the original (ground truth) data and the data generated by our model (both in-silico and interpolated slices), providing a clear metric for evaluating how well our model can replicate the complex spatial organization found in actual tissue samples.

#### Spatial Distribution of Normalized Spearman Correlation

The second evaluation metric used is the Spearman correlation, which assesses the similarity between ground truth gene expression data and interpolated gene expression data across spatial spots in a dataset. This approach is advantageous in spatial transcriptomics to understand how well the interpolation process preserves the original data’s spatial gene expression patterns.

For each spot *i*, we calculated the *ρ* spearman correlation coefficient between the corresponding rows (gene expression profiles) in the *X*_*ground truth*_ *and X*_*interpolated*_ as:

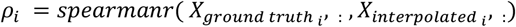

where:
*X*_*ground truth*_*and X*_*interpolated*_ denote the gene expression profiles for the spot *I* in the ground truth and interpolated data, respectively, and s*pearmanr*() is the function that computes the Spearman correlation coefficient. After calculating the Spearman correlation for each spot and comparing the original and interpolated expression matrices, these correlations are normalized. This normalization step adjusts the correlation values so they fit within a specific range, making it easier to visualize differences.

## References

1. Vandereyken, K., Sifrim, A., Thienpont, B. & Voet, T. Methods and applications for single-cell and spatial multi-omics. Nat Rev Genet 1–22 (2023) doi:10.1038/s41576-023-00580-2.

2. Zeng, Z., Li, Y., Li, Y. & Luo, Y. Statistical and machine learning methods for spatially resolved transcriptomics data analysis. Genome Biology vol. 23 Preprint at 10.1186/s13059-022-02653-7 (2022).

3. Fang, S. et al. Computational Approaches and Challenges in Spatial Transcriptomics. Genomics, Proteomics and Bioinformatics vol. 21 24–47 Preprint at 10.1016/j.gpb.2022.10.001 (2023).

4. Palla, G., Fischer, D. S., Regev, A. & Theis, F. J. Spatial components of molecular tissue biology. Nat Biotechnol 40, 308–318 (2022).

5. Zeira, R., Land, M., Strzalkowski, A. & Raphael, B. J. Alignment and integration of spatial transcriptomics data. Nat Methods 19, 567–575 (2022).

6. Wang, G. et al. Construction of a 3D whole organism spatial atlas by joint modelling of multiple slices with deep neural networks. Nat Mach Intell (2023) doi:10.1038/s42256-023-00734-1.

7. Ortiz, C. et al. Molecular Atlas of the Adult Mouse Brain. https://www.science.org (2020).

8. Ho, J., Jain, A. & Abbeel, P. Denoising diffusion probabilistic models. in Advances in Neural Information Processing Systems vols 2020-December (Neural information processing systems foundation, 2020).

9. Xiao, Z., Kreis, K. & Vahdat, A. TACKLING THE GENERATIVE LEARNING TRILEMMA WITH DENOISING DIFFUSION GANS. in ICLR (2022).

10. Yang, L. et al. Diffusion Models: A Comprehensive Survey of Methods and Applications. ACM Comput Surv 56, 1–39 (2024).

11. MoffiT, J. R. et al. Molecular, spatial, and functional single-cell profiling of the hypothalamic preoptic region. Science (1979) 362, (2018).

12. Palla, G. et al. Squidpy: a scalable framework for spatial omics analysis. Nat Methods 19, 171–178 (2022).

13. Maynard, K. R. et al. Transcriptome-scale spatial gene expression in the human dorsolateral prefrontal cortex. Nat Neurosci 24, 425–436 (2021).

14. Wang, X. et al. Three-dimensional intact-tissue sequencing of single-cell transcriptional states. Science (1979) 361, (2018).

15. Ho, J., Jain, A. & Abbeel, P. Denoising Diffusion Probabilistic Models. https://github.com/hojonathanho/diffusion.

16. Wang, H. et al. Scientific discovery in the age of artificial intelligence. Nature 620, 47–60 (2023).

17. Tegnér, J. N. et al. Computational disease modeling - Fact or fiction? BMC Syst Biol 3, (2009).

18. Zhang, J. et al. Spatiotemporal Omics-Refining the landscape of precision medicine. Life Medicine 1, 84–102 (2022).

19. Dong, K. & Zhang, S. Deciphering spatial domains from spatially resolved transcriptomics with an adaptive graph attention auto-encoder. Nat Commun 13, 1–12 (2022).

20. Wolf, F. A., Angerer, P. & Theis, F. J. SCANPY: Large-scale single-cell gene expression data analysis. Genome Biol 19, 15 (2018).

21. Loshchilov, I. & HuTer, F. Decoupled Weight Decay Regularization. 7th International Conference on Learning Representations, ICLR 2019 (2017).

