## Supplement Figures for "SpatialDiffusion: Predicting Spatial Transcriptomics with Denoising Diffusion Probabilistic Models"

**Supplementary Figure S1,** in silico generation of MERFISH data, **a)** showcases a heatmap of neighborhood enrichment, highlighting how stDiffusion effectively simulates spatial transcriptomics data in silico, ensuring the preservation of cluster neighborhoods as seen in actual data, **b)** presents a scatter plot of clusters identified through Leiden clustering and plotted in spatial coordinates, effectively reflecting the clustering and spatial distributions found in the actual data.

**Supplementary Figure S2,** spatial distribution of normalized Spearman correlation for DLPFC and Starmap data **a, b)** spatial distribution of normalized Spearman correlation between the ground truth Layer 4 from a DLPFC slice and interpolated Layer 4 with stDiffusion and linear interpolation, respectively, **c, d)** spatial distribution of normalized spearman correlation between the ground truth layer 2/3 from a Starmap slice and interpolated layer 2/3 with stDiffusion and linear interpolation. Results show that stDiffusion captures the spatial structure as it is distributed more evenly than linear interpolation, **e)** spatial distribution of normalized spearman correlation between the ground truth slice 9 from a mouse MERFISH data and interpolated slice 9 with stDiffusion, showing its efficacy in maintaining the spatial distribution of gene expression levels in the interpolated slice, consistent with the original data.

**Supplementary Figure S3, S4, S5**, Effect of lambda on interpolation within and across slices: Figures **S3 (DLPFC), S4 (Starmap), and S5 (MERFISH),** illustrate the interpolation results for varying lambda ($\lambda$) values from 0.1 to 0.9, demonstrating how different blending ratios between reference slices and layers within a slice affect the quality and accuracy of the interpolated gene expression profiles. Each subplot represents the interpolated data generated using a specific $\lambda$

value, highlighting the gradual transition in gene expression patterns as the blending ratio changes. The comparison underscores the significance of selecting an optimal λ value to achieve the most biologically plausible interpolation between slices.

Supplementary Figure S1

a)

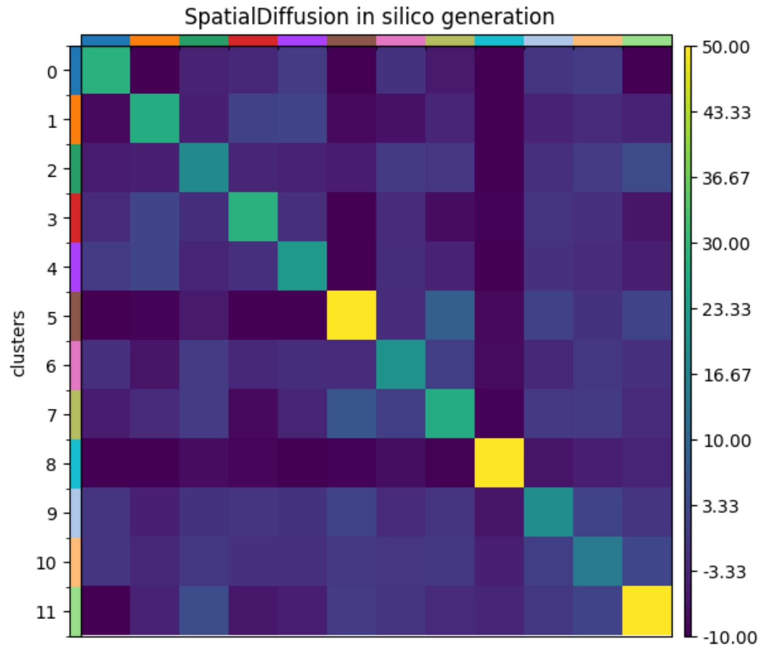


b)

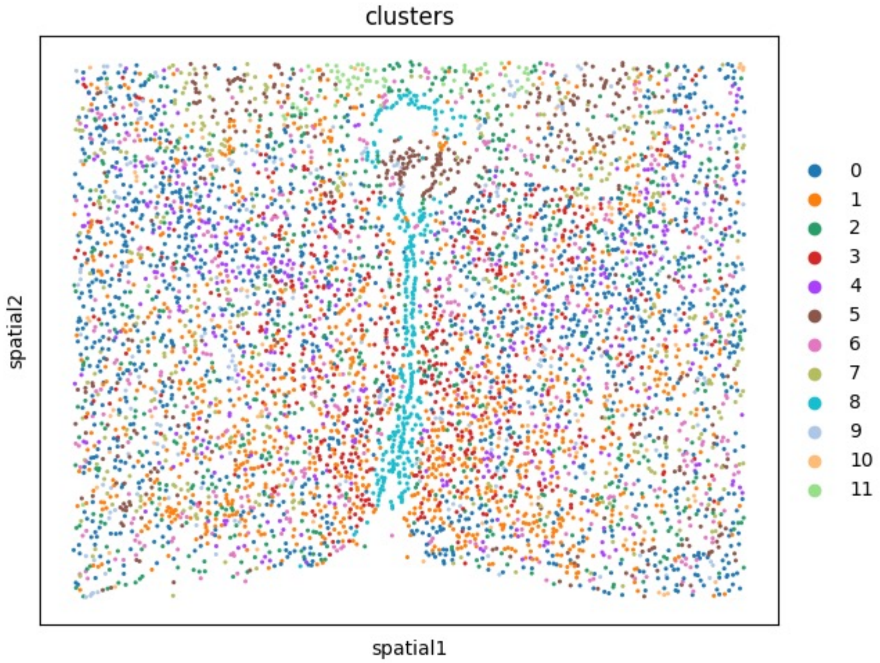

Supplementary Figure S2

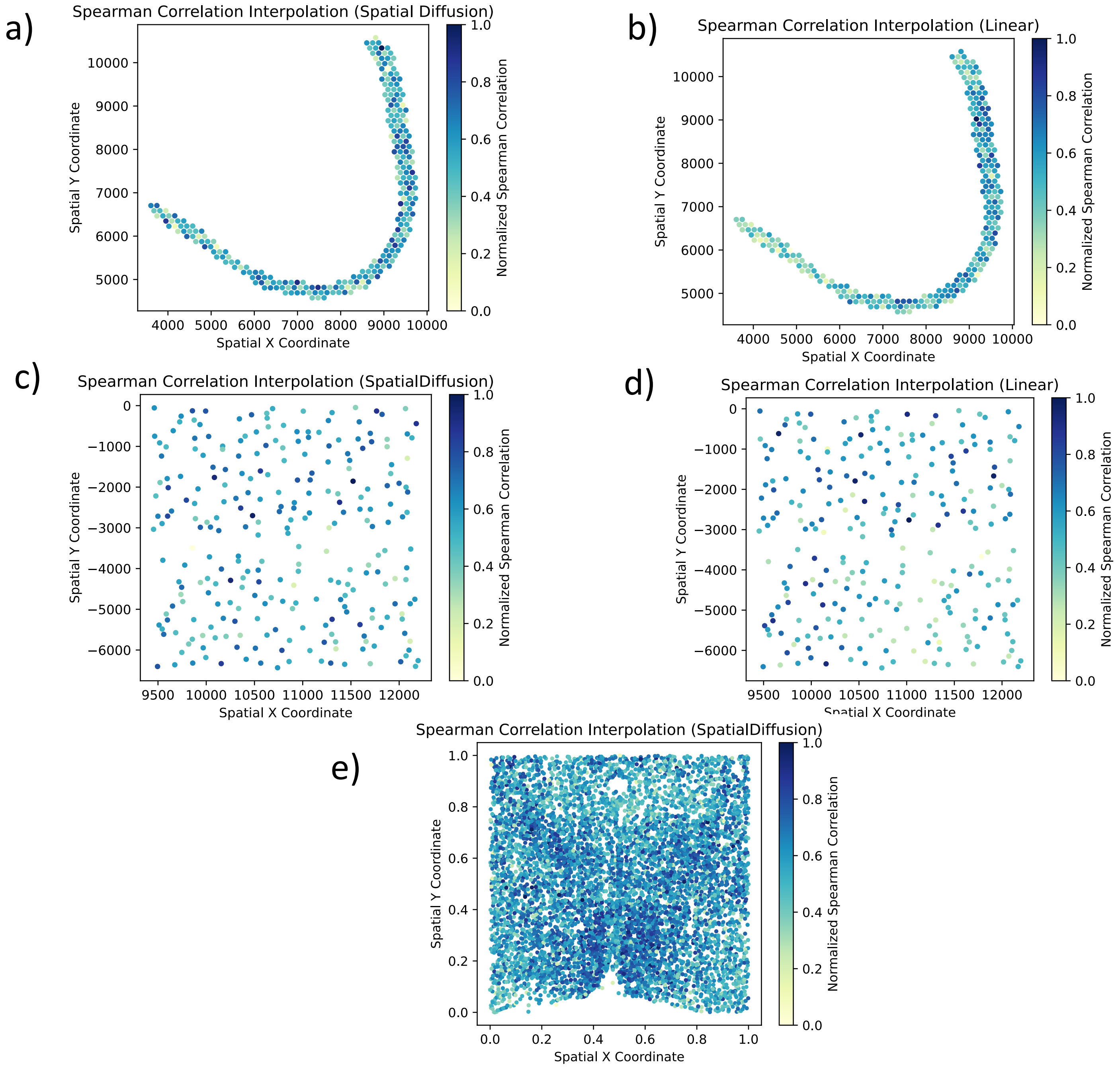

### Supplementary Figure S3

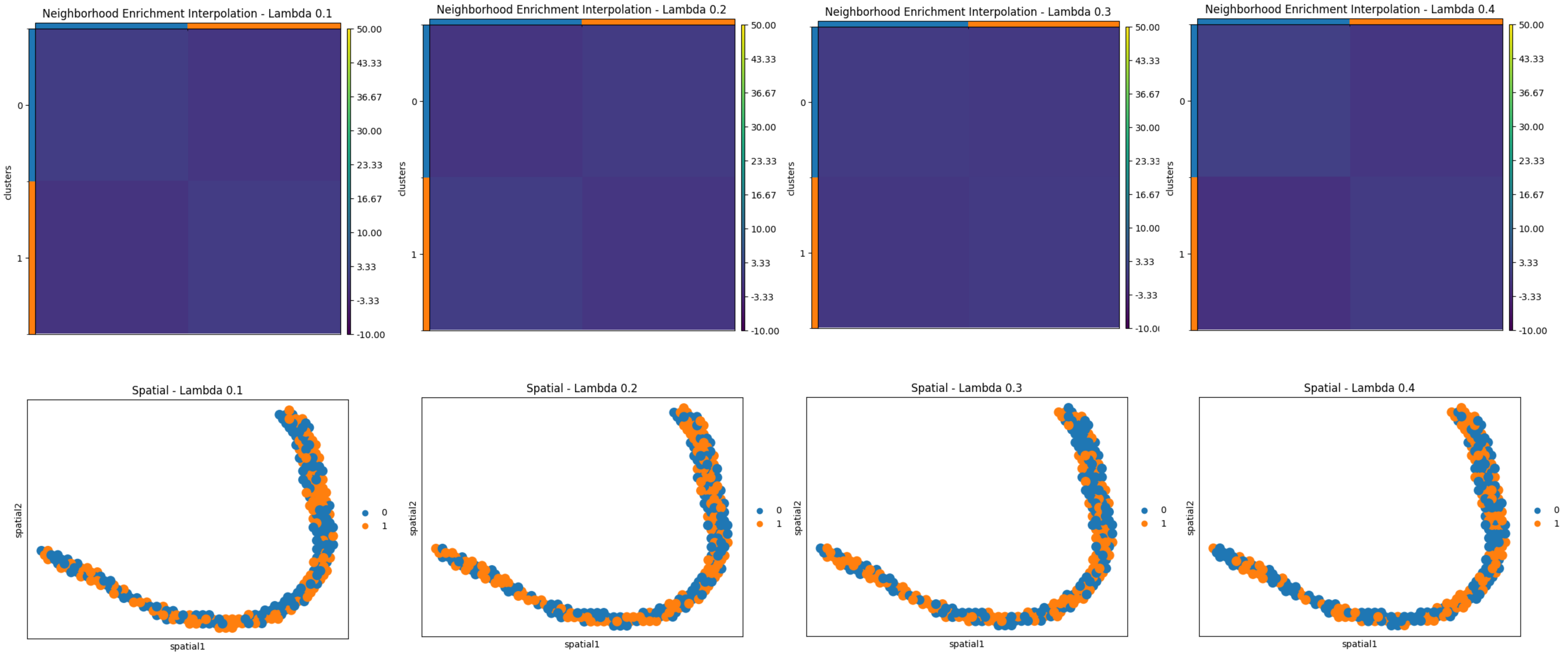

### Supplementary Figure S3

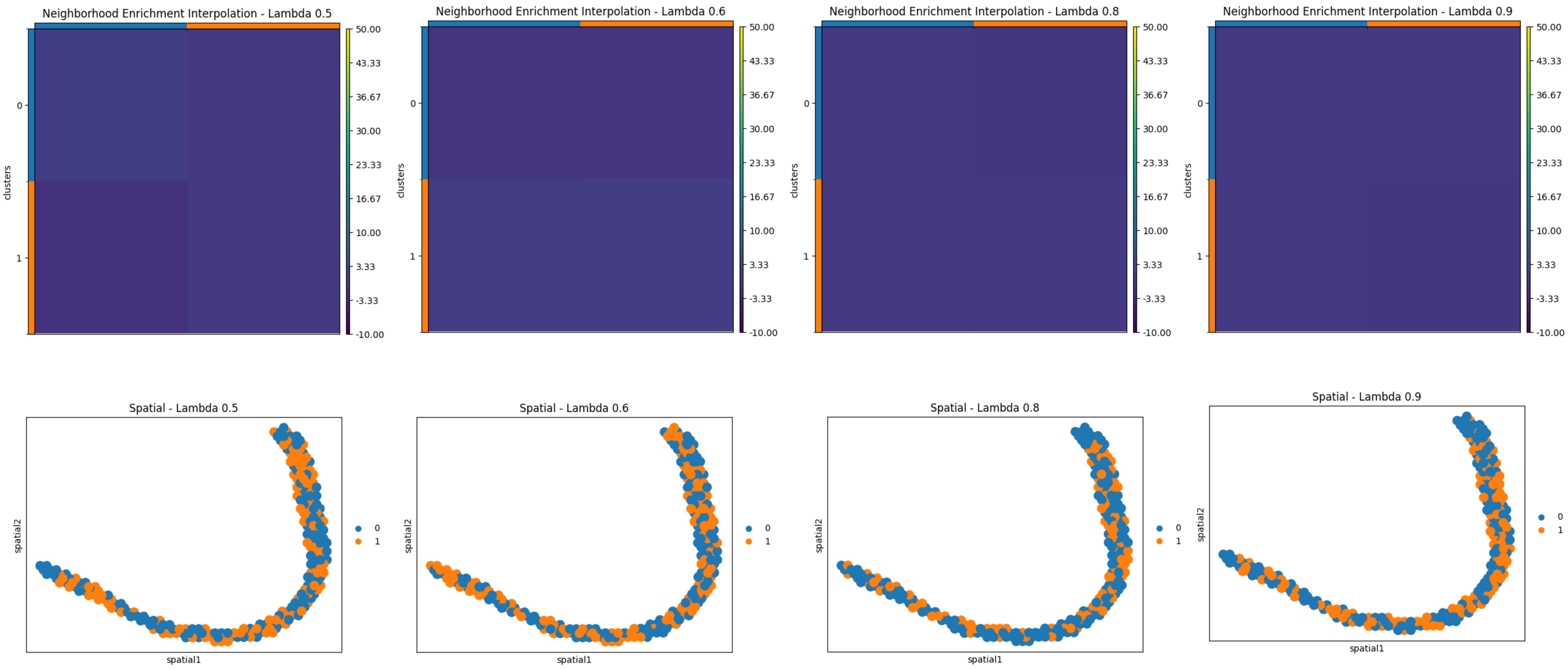

### Supplementary Figure S4

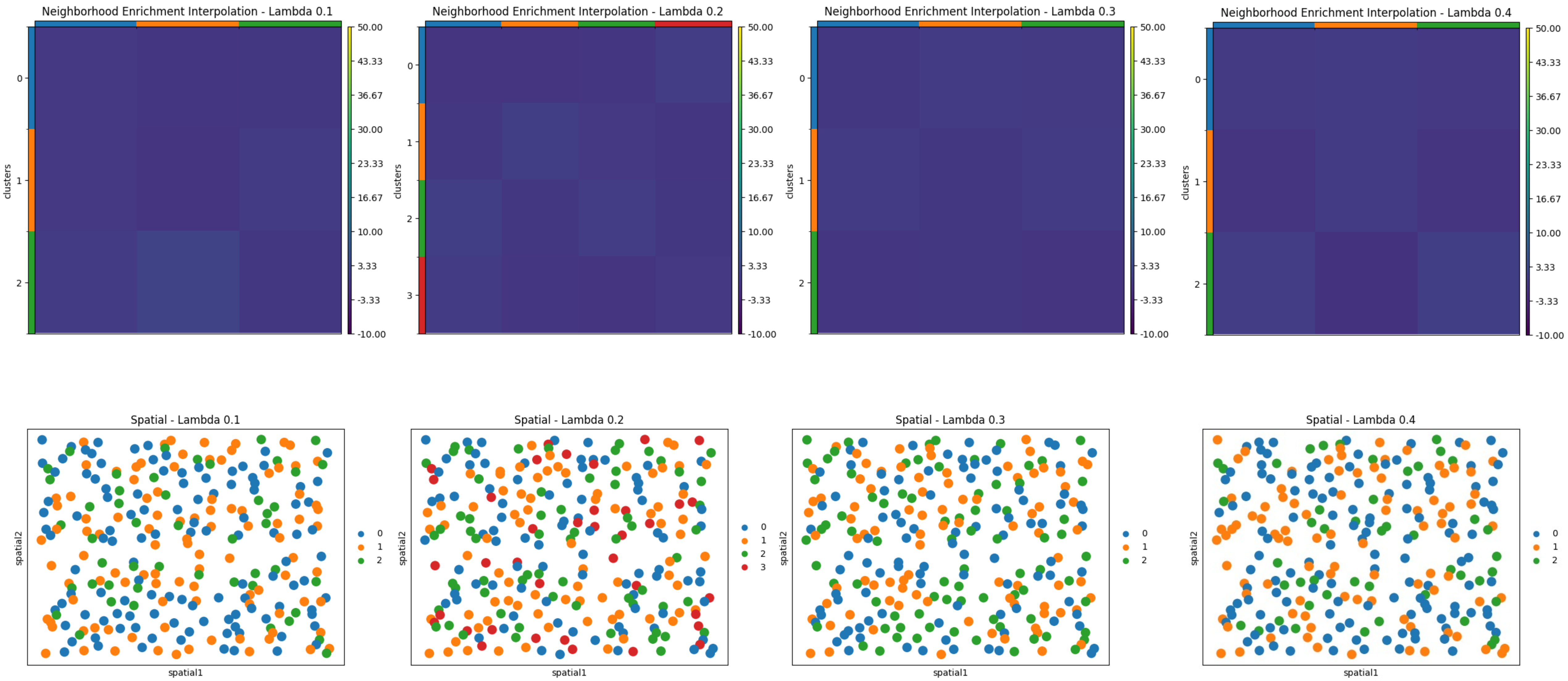

### Supplementary Figure S4

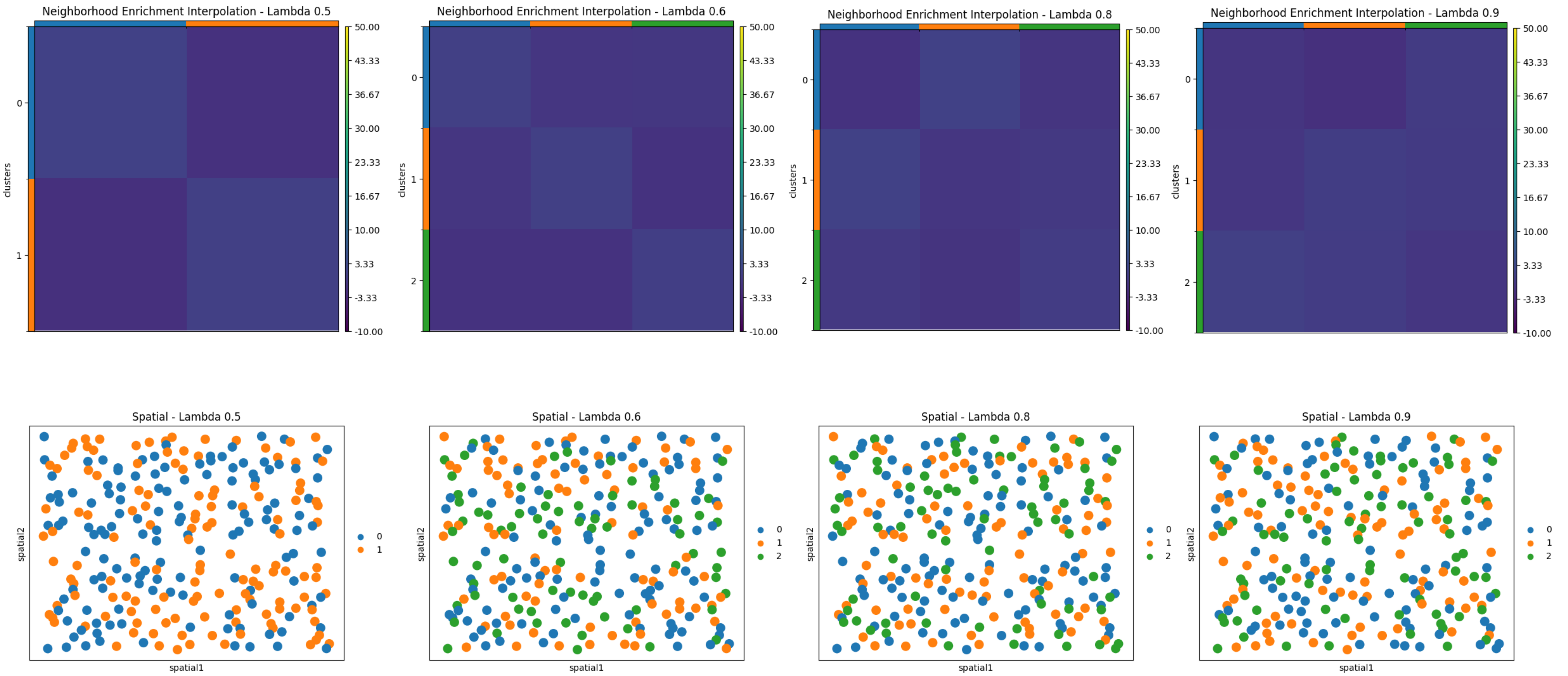

Supplementary Figure S5

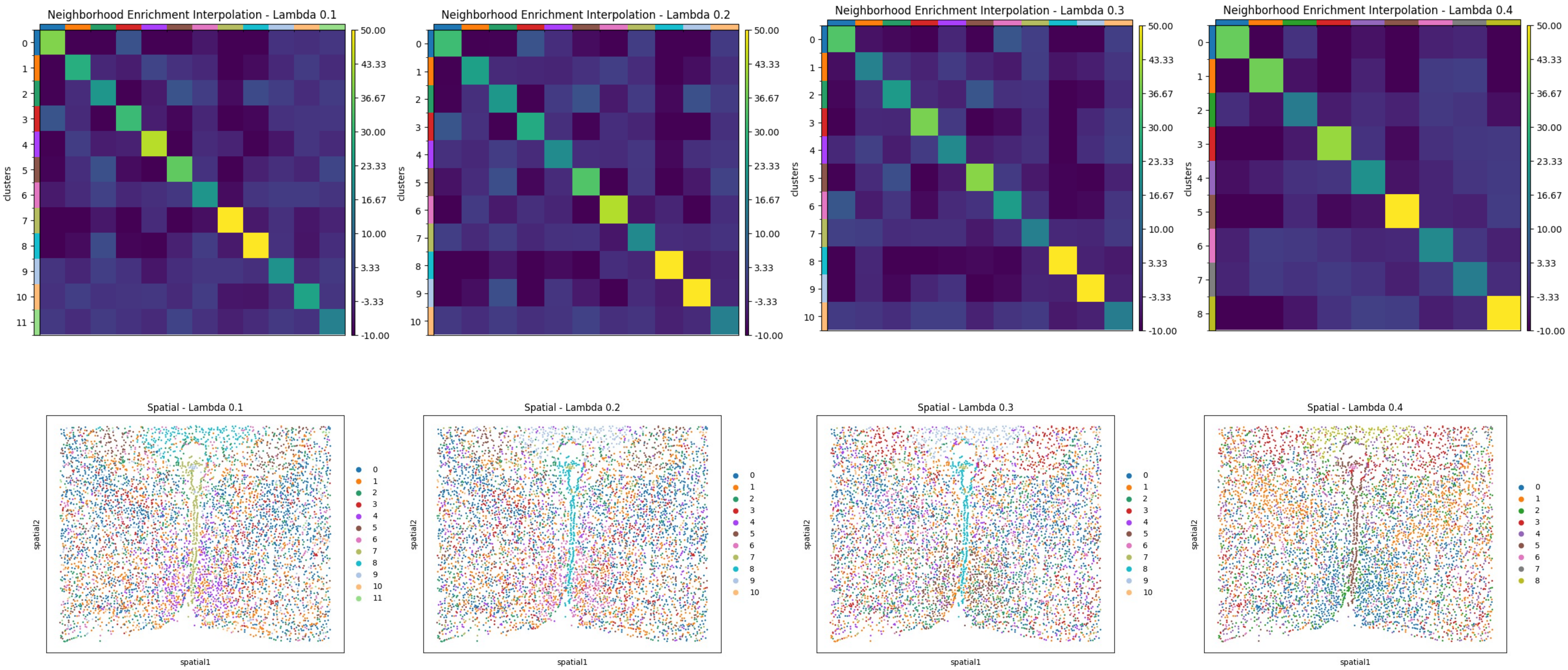

### Supplementary Figure S5

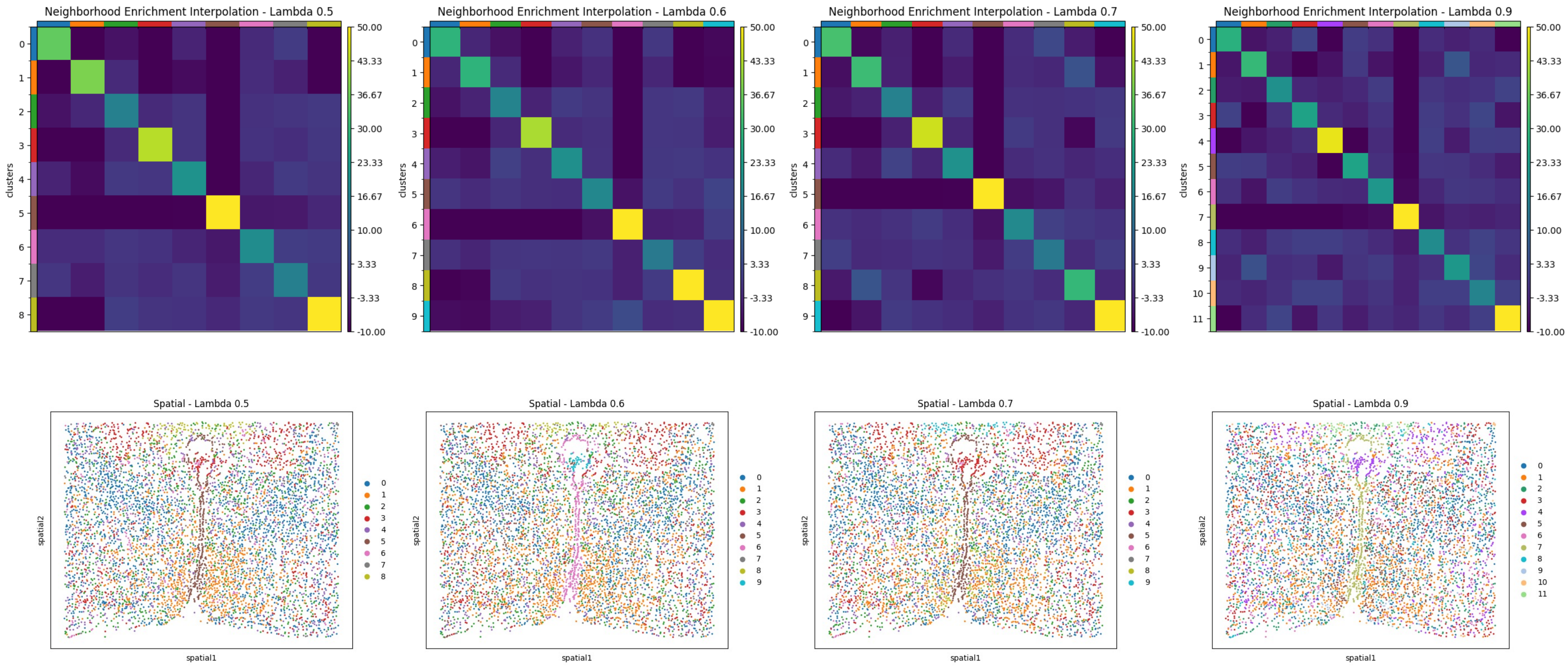
